# In-Solution Measurements of Binding Kinetics and Thermodynamics by nano Temperature Alteration Kinetics

**DOI:** 10.64898/2026.10.22.760345

**Authors:** Andreas Langer, Christian Osseforth, Philipp Schramm, Mariam Mohamadi, Nils-Eric Günther, Oscar Reinecke

## Abstract

Binding kinetics are central to mechanism-driven drug discovery. Previously, we established Spectral Shift technology for the measurement of equilibrium binding affinity by detecting small shifts in the emission spectrum of a fluorescently labeled molecule upon binding to a ligand. While slow binding kinetics can be observed as a gradual change toward equilibrium after mixing the binding partners, faster interactions often have already reached equilibrium once the measurement is started. Here, we extend Spectral Shift to the measurement of fast binding kinetics through a method termed “nano Temperature Alteration Kinetics” (nanoTAK). A rapid temperature step perturbs the binding equilibrium of an enthalpically driven interaction, and the subsequent relaxation toward the new equilibrium is monitored in real time by recording the time-dependent Spectral Shift signal. The relaxation directly encodes the association rate (*k*_on_) and dissociation rate (*k*_off_) of the interacting molecules, which are extracted by fitting data of a full ligand dilution series to a global kinetic model. In addition, the fit also returns thermodynamic parameters of the interaction such as the binding enthalpy and entropy. We validate the approach across three tiers of molecular interactions. First, hybridization of a 10-mer DNA duplex serves as a model system with large binding enthalpy. Global fits match the data over three different temperature steps and confirm the accuracy of the derived fit model. Second, the kinase p38α with a set of six ATP-competitive and allosteric inhibitors demonstrates that the method extends to a classical protein–small molecule system with substantially smaller enthalpic changes, yielding consistent rate constants across compounds and temperatures and offering unique insights into their temperature dependence. Third, the degrader system consisting of VHL E3-ligase complex (VCB), PROTAC MZ1 and bromodomain 2 of BRD4 (BRD4(BD2)) illustrates the application to a higher-order system, revealing differences in the binding kinetics between the binary and ternary complex. The presented method can resolve interactions that re-equilibrate faster than within ∼1 second, a timescale that can be difficult to access reliably with surface-based instruments. Binding kinetics can be measured both near room temperature, using a temperature step from 20 to 25 °C, but also, as an inherent feature of the approach, at physiologically relevant temperatures such as 37 °C.

## Introduction

The equilibrium dissociation constant (*K*_d_) is a key descriptor for the strength of a molecular interaction, yet it does not capture the lifetime of a molecular complex. Two ligands can share an identical *K*_d_ but behave very differently in a cell if one dissociates in seconds and the other in hours. Residence time, and the underlying rate constants *k*_on_ and *k*_off_ that define it, therefore provide an important additional dimension for interaction characterization in drug discovery^1,2^.

Surface plasmon resonance (SPR), the established workhorse for kinetic measurements, immobilizes one binding partner on a sensor surface while the other flows across it. Biolayer interferometry (BLI) follows the same basic principle: one partner is immobilized on a sensor tip that is then dipped into a solution containing the other. This surface-based architecture is powerful, but it also imposes inherent constraints. Immobilization can alter the behavior of the captured partner by blocking access to its binding site, restricting its conformational freedom, or reducing its activity^3,4^. Certain target classes, including membrane proteins and large multiprotein assemblies, are especially difficult to study in this format. Even in well-behaved systems, surface-based methods are vulnerable to mass transport limitation and analyte rebinding, both of which can skew apparent rate constants and require considerable expertise to detect and control^3^. Fast interactions pose a further challenge: when re-equilibration occurs faster than surface mixing, the recorded sensorgram reflects the limitations of fluidics rather than the underlying binding kinetics.

A solution-phase method that requires no immobilization would circumvent these limitations, but it introduces a different challenge: how to perturb an equilibrium that is, by design, already at rest. Mixing-based relaxation, as implemented in stopped-flow experiments, is one approach; thermal perturbation is another. Classical temperature-jump relaxation kinetics, pioneered by Eigen and co-workers, takes advantage of the temperature dependence of equilibrium in enthalpy-driven interactions^5^. A rapid temperature step displaces the system from equilibrium, and the subsequent relaxation to the new equilibrium reports directly on the underlying kinetic rate constants. Although the theory is well established, classical temperature-jump studies of biomolecular systems often required specialized instrumentation and relatively large, concentrated samples to produce measurable relaxation signals^6,7^.

Here we implement temperature-jump relaxation in a microliter-scale, capillary-based format with a fluorescence readout, removing both the immobilization requirement of surface methods and the sample burden of classical T-Jump measurements, a new method we term “nano Temperature Alteration Kinetics” (nanoTAK). A fluorescently labeled target and a titrated binding partner are prepared exactly as for a conventional solution affinity measurement and then loaded into glass capillaries. A collimated infrared laser is modulated in a way so that it creates a rapid and spatially homogeneous temperature step of defined magnitude within approximately 250 milliseconds. The resulting relaxation of the system is monitored in real time through measurement of the Spectral Shift signal of the labeled partner (the 670 nm / 650 nm fluorescence emission ratio), which reports on the changes in the bound fraction^8^. Because the same sample is measured at the same position throughout, the approach avoids capillary-to-capillary variation and enables detection of even smallest signal changes. Kinetic rate constants are then obtained by fitting the data from a full dilution series to a global in-solution binding model.

We validate the method across three systems of increasing complexity. First, the hybridization of a short DNA duplex serves as benchmark: the binding partners are thermally stable, and the binding enthalpy of the interaction is roughly tenfold larger than a typical protein–small molecule interaction, producing an exceptionally large and clean relaxation signal. Having validated the fitting model, we next investigate the kinase p38α with a set of six ATP-competitive and allosteric inhibitors. It is a system where the enthalpic changes are far smaller and the equilibrium moves much less. Finally, we assess binding of the PROTAC MZ1 to the VHL E3 ligase complex (VCB) in absence and presence of the bromodomain 2 of BRD4 (BRD4(BD2)). It is a well-characterized system that has previously been extensively studied using isothermal titration calorimetry (ITC) and SPR and thus allows the comparison of the derived kinetic and thermodynamic parameters with those from established methods. The system presents an additional challenge because of its rapid binding kinetics and the small enthalpic contribution within the binary complex, as well as the involvement of multiple protein complexes in the ternary interaction, since VCB is itself a multisubunit complex.

## Materials & Methods

### nanoTAK measurements

All nanoTAK measurements in this work were performed in coated glass capillaries (cat# MO-K025, NanoTemper Technologies GmbH) using the Monolith Omni instrument (NanoTemper Technologies GmbH) equipped with dual-emission Spectral Shift detection optics and a collimated IR-laser. The collimated laser heats the capillary observation volume homogeneously and can thereby produce a rapid, well-defined temperature step. The required volume per data point was 10 µL. Environment-sensitive red dyes with an emission maximum at around 660 nm (e.g., RED 2nd Generation dyes, NanoTemper Technologies GmbH, or Cy5) were used as fluorescent reporters.

For all interaction measurements, the fluorescently labeled target was held at constant concentration while the binding partner was titrated across a broad range spanning baseline to saturation (12 or preferably 16 different concentrations). Samples were loaded into glass capillaries and equilibrated at the set capillary tray temperature for at least 5 minutes before the nanoTAK measurement (20, 22 or 25 °C). If not stated otherwise, three technical replicates per temperature jump were performed on the same set of capillaries. Re-establishment of the initial equilibrium served as a quality control measure for reversibility. For each system, temperature jumps of +5, +10, and +15 °C were applied.

Relaxation to the new equilibrium was monitored by recording the Spectral Shift ratio (670 nm / 650 nm) of the fluorescently labeled partner at 50 ms sampling intervals. The acquisition time per sample ranged from 5 s for fast binding kinetics up to 60 s for slower interactions.

### IR-laser calibration

The required IR-laser modulation for each temperature step was calibrated using a photostable red fluorescent dye with a reproducible temperature-dependent fluorescence intensity. Reference data were recorded in an Andromeda X device (NanoTemper Technologies GmbH) to create a fluorescence–temperature calibration curve. The laser-power modulation array (updated every 250 ms) was iteratively refined so that the median normalized fluorescence of a set of dye-filled capillaries reached and held the target value corresponding to the desired temperature jump. Jumps of +5, +10, and +15 °C were used during the calibration. For all temperature jumps, the target plateau was reached within the first 250 ms.

### DNA hybridization

Chemically synthesized DNA oligonucleotides were obtained from metabion international AG (Planegg, Germany). A fluorescently labeled 10-mer oligonucleotide (sequence 5’ Cy5 GGA CTT CAG G 3’) was designed as target strand and the complementary 10-mer (sequence 3’ CCT GAA GTC C 5’) was used as ligand strand.

Lyophilized oligonucleotides were resuspended in double-distilled water (ddH_2_O) to a concentration of 100 µM. Assay buffer consisting of 0.25X PBS buffer supplemented with 0.005 % TWEEN® 20 was prepared as a two-fold stock solution. The target strand was further diluted in two-fold assay buffer, while the ligand strand was serially diluted in ddH_2_O. The highest final concentration of the ligand strand in the 16-point dilution series was 5 µM. nanoTAK measurements were performed at 25 °C using temperature jumps of +5, +10, and +15 °C. The duration of each measurement was 15 seconds.

### Protein labeling

Recombinant p38α, VCB (the complex of VHL E3 ligase, Elongin C and Elongin B) and BRD4(BD2) were obtained from CRELUX, a WuXi AppTec Company (Munich, Germany).

Covalent labeling of cysteine residues of p38α was performed using the Spectral Shift Optimized Protein Labeling Kit - Cysteine-Reactive (cat# MO-L024, NanoTemper Technologies GmbH). In brief, p38α was prepared at a final concentration of 20 µM and a three-fold molar excess of dye was added, followed by incubation for 30 minutes at room temperature in the dark. Unreacted dye was removed using the B-Column of the labeling kit. Protein concentration and degree-of-labeling were measured afterwards with a NanoPhotometer NP80 photo-spectrometer (Implen, Munich, Germany).

VCB was labeled with the Protein Labeling Kit RED-Maleimide 2nd Generation (cat# MO-L014, NanoTemper Technologies GmbH) using the same procedure described above.

### Protein–small molecule interactions

The p38α inhibitors SB203580, SB202190, SB239063, RWJ67657, PD169316, and BIRB796 were purchased from Sigma-Aldrich and prepared as 100 µM stock solutions in DMSO. Working solutions were prepared by mixing 2 µL of each 100 µM inhibitor stock with 98 µL of assay buffer containing 20 mM HEPES (pH 7.4), 150 mM NaCl, 2 mM GSH, 1 mM EDTA, and 0.005 % TWEEN® 20. All subsequent dilutions were performed in assay buffer supplemented with 2 % DMSO.

BIRB796 was tested up to a maximum concentration of 300 nM, while the final highest concentration of all other inhibitors was 1 µM. The final concentration of labeled p38α in all assays was 5 nM, and the final DMSO concentration was maintained at 2 %. nanoTAK measurements were performed at 20 °C using temperature jumps of +5, +10, and +15 °C. The duration of each nanoTAK measurement was 10 seconds.

### Binary and ternary complex interactions

The PROTAC MZ1 was ordered from opnMe (Ingelheim, Germany). All measurements were performed in assay buffer containing 50 mM Tris (pH 7.8), 150 mM NaCl, 10 mM MgCl_2_, 2 mM GSH, and 0.01 % Pluronic® F-127, with a final DMSO concentration of 2 %.

For the binary measurements, the final concentration of labeled VCB was 10 nM, and MZ1 was tested up to a maximum concentration of 2 µM. For the ternary complex measurements, the final concentration of labeled VCB was 2.5 nM, MZ1 was tested up to 200 nM, and BRD4(BD2) was maintained at a constant concentration of 400 nM in all samples. MZ1 was prepared as a 12-point, twofold serial dilution.

nanoTAK measurements were performed at 20 °C using temperature jumps of +5, +10, and +15 °C. The measurement duration was 5 s when characterizing the binary complex and 60 s for the ternary complex.

### Data analysis

A conventional Spectral Shift affinity measurement at the set capillary tray temperature *T*_0_ provides the initial dissociation constant *K*_d_(*T*_0_) before the temperature jump and fixes the starting bound fraction *f*_0_ at each concentration according to the standard equation for 1:1 binding:

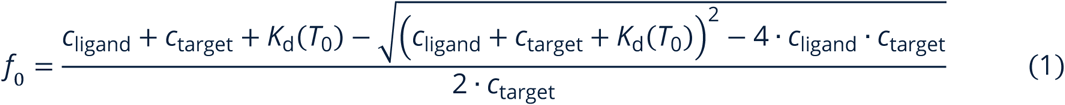

where *c*_ligand_ and *c*_target_ are the total ligand and target concentrations within the sample.

Time-resolved relaxation data for the full series is then globally fitted to an in-solution kinetic model, returning *k*_on_, *K*_d_ and *k*_off_ = *k*_on_ · *K*_d_ at the elevated second temperature *T*_1_ after the temperature jump.

The fraction of bound target *f* (*t*) is described by:

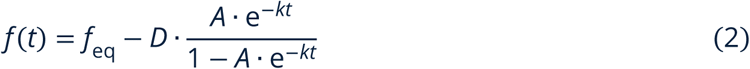

where

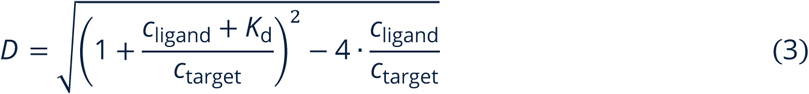

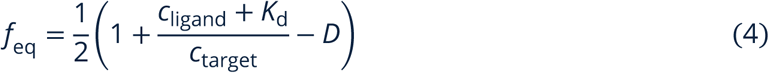

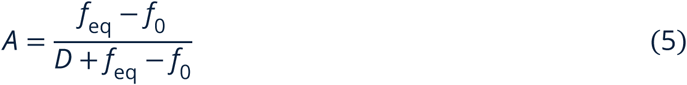

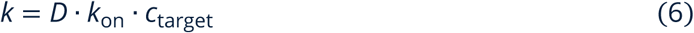

A detailed derivation of Eq. (2) is available in the Supplementary Information. The time-resolved Spectral Shift response is then fitted by the equation

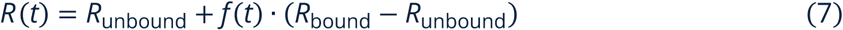

where *R* (*t*) is the 670 nm / 650 nm ratio trace at a given ligand concentration *c*_ligand_. *R*_unbound_ and *R*_bound_ are the equilibrium ratio values of the target alone and complex at the elevated temperature. In the presence of ligand-induced changes of the fluorescence intensity at 650 nm, a mathematical correction is applied as described previously^8^.

To derive the van’t Hoff binding enthalpy, we assume that it is approximately constant between *T*_0_ and *T*_1_, and then estimate it from the two-point van’t Hoff relationship:

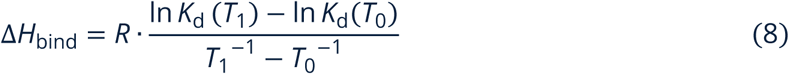

where *R* is the ideal gas constant.

The entropic contribution is then calculated using the Gibbs relation at *T*_0_:

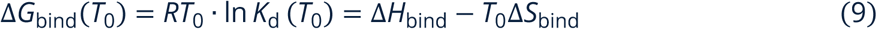

## Results

### Method principle and kinetic model

In a nanoTAK experiment, each capillary contains a labeled target at fixed concentration and a binding partner at one concentration of a dilution series, pre-equilibrated at the starting temperature *T*_0_ (typically 20 or 25 °C). A power-modulated collimated IR-laser drives a rapid step to a temperature *T*_1_ > *T*_0_. The temperature reaches a flat plateau within approximately 250 ms and is maintained within ± 0.25 °C of the defined target temperature *T*_1_ for the duration of the measurement (Fig. 1A). A stable plateau is essential, since rate constants are themselves temperature-dependent and must be read at a single, well-defined temperature. The step-like temperature change is achieved by an initial high IR-laser pulse which then is continuously reduced to a constant, non-zero power value to maintain the final desired temperature (Fig. 1B).

For enthalpically driven interactions, the equilibrium dissociation constant at *T*_1_ differs from its value at *T*_0_, with the magnitude and direction of the change determined by the van’t Hoff relationship and hence by the sign and magnitude of the binding enthalpy. The system, momentarily out of equilibrium, relaxes toward the new equilibrium via association and dissociation governed by *k*_on_ and *k*_off_ at *T*_1_.

Rate constants are obtained by globally fitting the complete dilution series data to an in-solution kinetic model. For each concentration, the model starts from a known, nonzero bound fraction that is calculated from the *K*_d_ measured at *T*_0_ in a conventional Spectral Shift binding affinity measurement performed on the same samples immediately before the IR-laser perturbation. The model also accounts explicitly for depletion of both partners, rather than assuming analyte excess as in pseudo-first-order surface analyses.

Importantly, the information needed to constrain the fit is taken from the full concentration series. A single relaxation curve cannot reliably resolve the multiple fit parameters, and we recommend at least 8 concentrations, with 12–16 preferred. The fit yields *k*_on_, *k*_off_, and *K*_d_ at the elevated temperature *T*_1_.

**Fig. 1:**
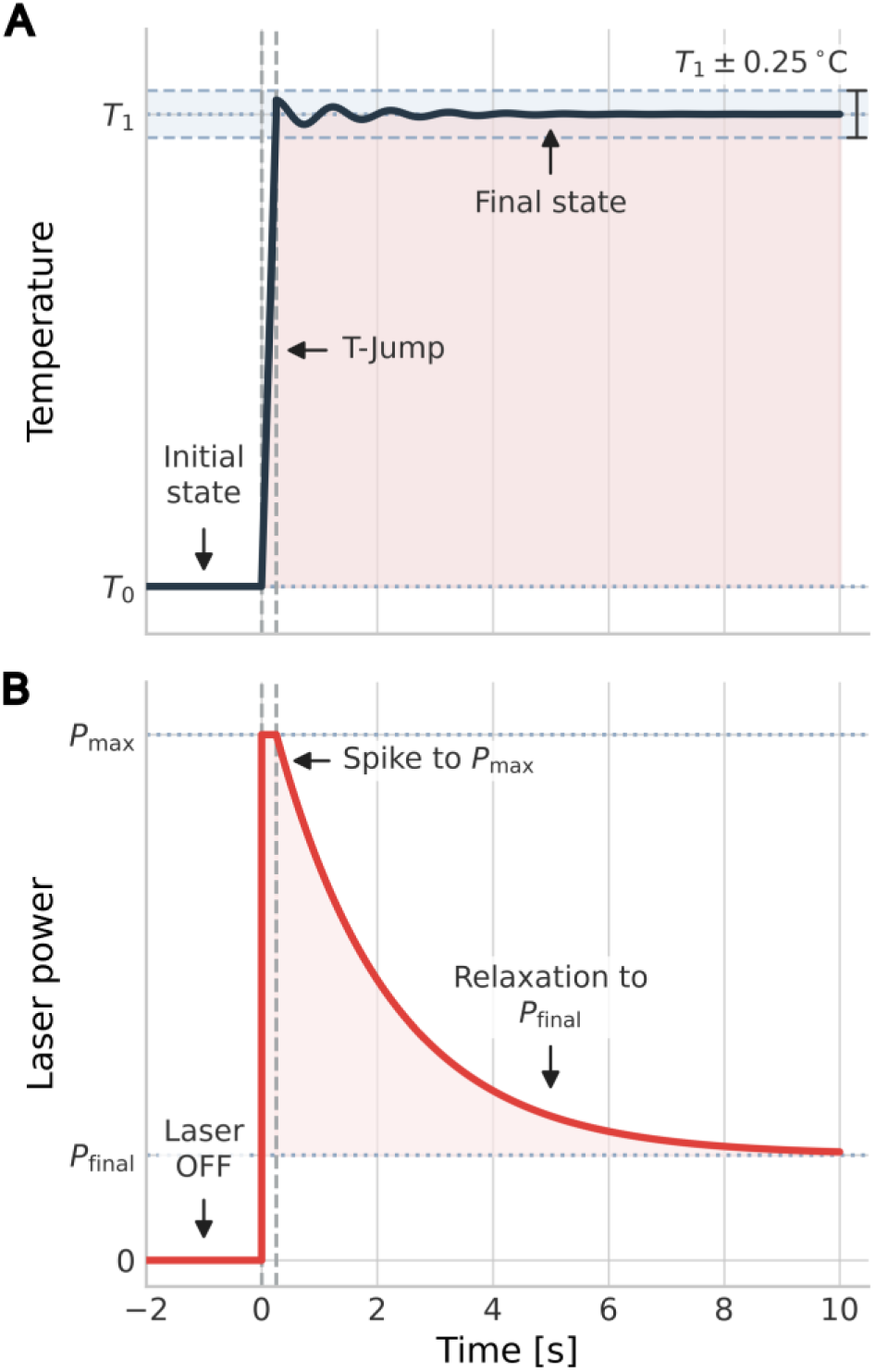
Temperature-jump profile and corresponding IR-laser power modulation for a nanoTAK measurement. (**A**) Schematic temperature profile during a nanoTAK experiment. The sample, initially equilibrated at *T*_0_, is rapidly perturbed to *T*_1_ within approximately 250 ms. Afterwards, the elevated temperature is maintained within ± 0.25 °C of the defined target temperature *T*_1_. (**B**) The IR-laser power profile used to implement the perturbation. A brief initial pulse at *t* = 0 with laser power *P*_max_ drives the rapid temperature increase, followed by relaxation to a lower steady-state power *P*_final_ that maintains the final temperature *T*_1_ within the capillary.

### DNA hybridization

DNA hybridization is an almost ideal model system for nanoTAK. Short DNA duplexes melt and reanneal reversibly and there is minimal risk of irreversibly denaturing one of the binding partners, as can be the case for a thermally unstable protein. Duplex formation is strongly enthalpically favorable, with a calculated enthalpy of approximately −30 kJ mol^−1^ per nearest-neighbor base-pair step^9^. For a 10-mer duplex, this corresponds to an interaction enthalpy substantially larger than that of a typical enthalpically driven protein–small-molecule interaction. The large absolute value of Δ*H* means the equilibrium *K*_d_ shifts strongly with temperature; or in other words, the DNA duplex melts when the temperature is increased. The large relaxation signal can therefore serve as a test for the method and validation of the kinetic model to extract kinetic and thermodynamic parameters.

We measured hybridization of a 10-mer duplex with one strand carrying a Cy5 label. The capillary tray was equilibrated at 20, 22 or 25 °C, and relaxation was recorded after temperature jumps of +5, +10, or +15 °C, with three replicates per condition. Excellent relaxation data were obtained for all three jump amplitudes, and the global in-solution model fit the experimental traces well across the full dilution series at each starting temperature. The 22 °C dataset is shown in Fig. 2; the corresponding datasets obtained at 20 and 25 °C are provided in Figs. S1 and S2.

The derived rate constants reveal an asymmetric temperature dependence that is characteristic of DNA duplex hybridization: the association rate *k*_on_ remains slow and essentially constant across the three temperature jumps (ranging from 2.3 to 3.4 × 10^5^ M^−1^s^−1^), while the dissociation rate *k*_off_ increases dramatically with temperature, rising roughly 100-fold from 0.016 s^−1^ at 25 °C to 1.9 s^−1^ at 40 °C (Table 1). It is this steep rise in *k*_off_, rather than any change in *k*_on_, that drives the strong decrease in binding affinity with increasing temperature. This behavior is consistent with established kinetic measurements of short DNA duplexes: literature values for DNA hybridization on-rates typically cluster around 10^5^–10^6^ M^−1^s^−1^ and show little dependence on temperature, whereas off-rates span many orders of magnitude and are strongly temperature-dependent, reflecting the large activation energy for strand melting^10,11^. The van’t Hoff binding enthalpy derived at each temperature jump is consistently large and negative, with values of around −230 to −260 kJ mol^−1^ (Table 1).

**Fig. 2:**
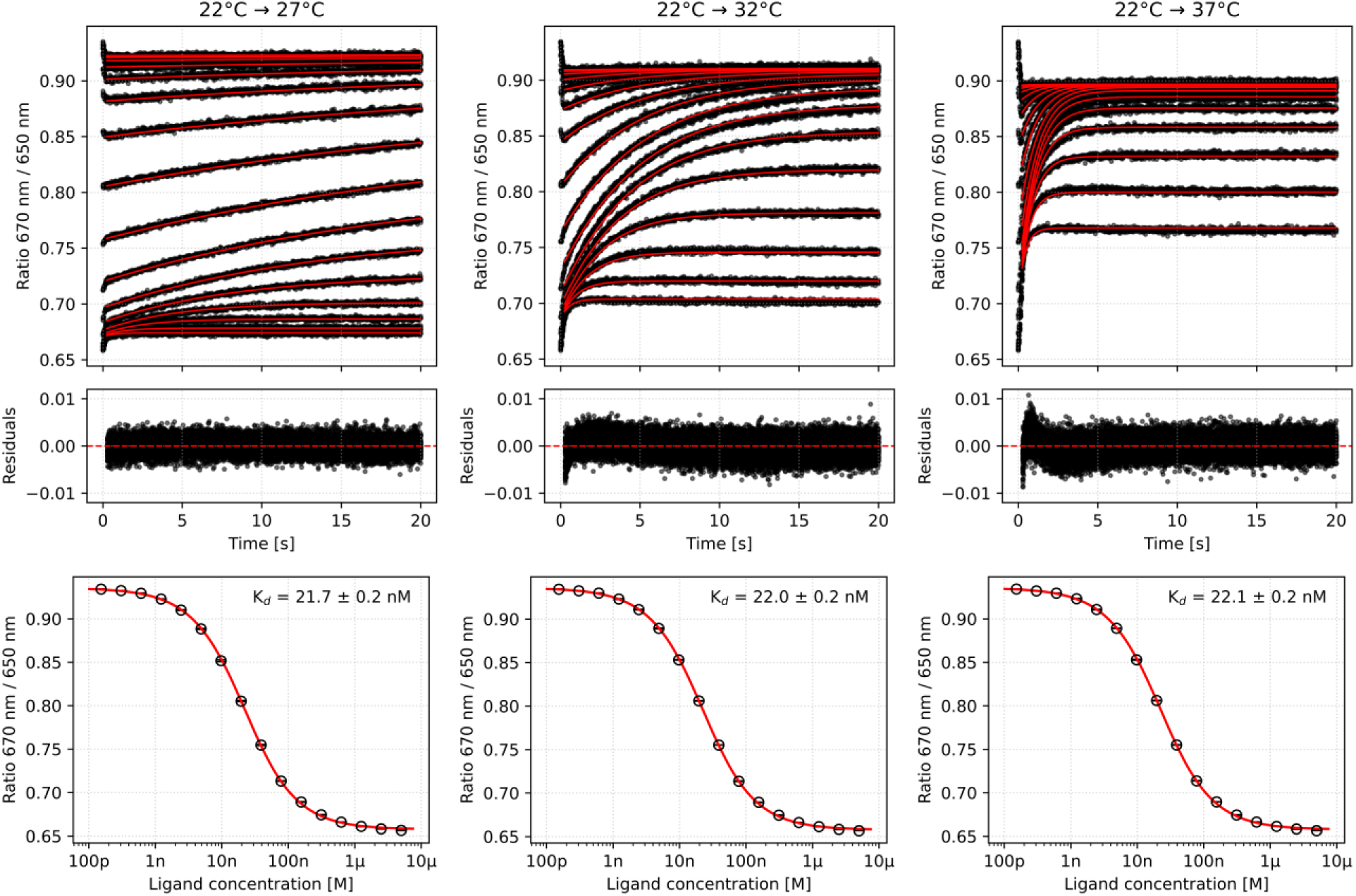
nanoTAK measurements of DNA hybridization. A 10-mer Cy5-labeled strand mixed with a serial dilution series of complementary unlabeled strand was measured at an initial capillary temperature of 22 °C. **Top row:** Experimental relaxation traces after temperature jumps of +5, +10, and +15 °C from left to right (black) and corresponding global in-solution model fits (red). The traces represent triplicate measurements across the dilution series. **Middle row:** Residuals of the global fits. **Bottom row:** Equilibrium dose response data measured before each temperature jump, together with the corresponding fits. The fitted initial *K*_d_ values were consistent across the three measurements, supporting reproducible re-equilibration of the system before each temperature-jump experiment. Equivalent datasets acquired at initial capillary temperatures of 20 and 25 °C are shown in Figs. S1 and S2.

The experimental design also enables the same kinetic parameters to be determined through multiple independent measurements. For example, *k*_on_ at 30 °C can be obtained from temperature jumps starting at either 20 or 25 °C. Likewise, *K*_d_ at 25 °C can be determined kinetically from a 20 to 25 °C jump and independently by an isothermal Spectral Shift measurement at 25 °C. The close agreement between these values, summarized in Table 1, validates the global in-solution analysis and supports the robustness of the derived binding parameters.

**Table 1:** Kinetic and thermodynamic parameters from nanoTAK measurements of DNA (de-)hybridization.

| Method | $T$ (°C) | $K_d$ (nM) | $k_{on}$ ($10^5 \text{ M}^{-1}\text{s}^{-1}$) | $k_{off}$ ($\text{s}^{-1}$) | $\Delta G$ ($\text{kJ} \times \text{mol}^{-1}$) | $\Delta H$ ($\text{kJ} \times \text{mol}^{-1}$) | $-T\Delta S$ ($\text{kJ} \times \text{mol}^{-1}$) |
| --- | --- | --- | --- | --- | --- | --- | --- |
| SpS | 20 | $11.3 \pm 0.2$ | — | — | — | — | — |
| nanoTAK | 25 | $55.8 \pm 0.7$ | $2.8 \pm 0.1$ | $0.016 \pm 0.001$ | $-44.6 \pm 0.1$ | $-232.2 \pm 2.5$ | $187.6 \pm 2.5$ |
| nanoTAK | 30 | $299 \pm 5$ | $3.1 \pm 0.1$ | $0.092 \pm 0.001$ | $-44.6 \pm 0.1$ | $-242.5 \pm 1.5$ | $197.8 \pm 1.5$ |
| nanoTAK | 35 | $1580 \pm 20$ | $3.3 \pm 0.1$ | $0.52 \pm 0.01$ | $-44.6 \pm 0.1$ | $-246.9 \pm 0.8$ | $202.3 \pm 0.8$ |
| SpS | 22 | $21.7 \pm 0.2$ | — | — | — | — | — |
| nanoTAK | 27 | $109 \pm 1$ | $2.9 \pm 0.1$ | $0.031 \pm 0.001$ | $-43.3 \pm 0.1$ | $-237.8 \pm 1.9$ | $194.5 \pm 1.9$ |
| nanoTAK | 32 | $571 \pm 5$ | $3.4 \pm 0.1$ | $0.19 \pm 0.01$ | $-43.3 \pm 0.1$ | $-243.8 \pm 0.8$ | $200.6 \pm 0.8$ |
| nanoTAK | 37 | $3090 \pm 30$ | $3.0 \pm 0.1$ | $0.92 \pm 0.01$ | $-43.3 \pm 0.1$ | $-250.6 \pm 0.6$ | $207.4 \pm 0.6$ |
| SpS | 25 | $58.3 \pm 0.5$ | — | — | — | — | — |
| nanoTAK | 30 | $291 \pm 3$ | $3.2 \pm 0.1$ | $0.093 \pm 0.001$ | $-41.3 \pm 0.1$ | $-242.2 \pm 1.6$ | $200.9 \pm 1.6$ |
| nanoTAK | 35 | $1600 \pm 20$ | $3.3 \pm 0.1$ | $0.53 \pm 0.01$ | $-41.3 \pm 0.1$ | $-252.9 \pm 0.8$ | $211.6 \pm 0.8$ |
| nanoTAK | 40 | $8290 \pm 70$ | $2.3 \pm 0.1$ | $1.9 \pm 0.1$ | $-41.3 \pm 0.1$ | $-256.2 \pm 0.5$ | $214.9 \pm 0.5$ |

Kinetics and thermodynamic parameters obtained from nanoTAK measurements for the hybridization between two 10-mer DNA strands. Kinetic data were acquired by triplicate measurements on the same set of capillaries. SpS: Equilibrium Spectral Shift measurement. $\Delta G$ and $-T\Delta S$ are reported for the initial sample temperature (the temperature of the equilibrium SpS measurement), i.e. 20 °C, 22 °C and 25 °C respectively.

### Protein–small molecule interactions

Having validated the model on a high-enthalpy system, we asked whether the method can also determine binding kinetics of systems where the equilibrium moves to a comparatively much lower extent. Protein–small molecule interactions typically have far smaller binding enthalpies, so the temperature-induced *K*_d_ shift and therefore the amplitude of relaxation is much smaller. To investigate this, we performed nanoTAK measurements of the mitogen-activated protein kinase p38α with a panel of well-characterized small molecule inhibitors (SB203580, SB202190, SB239063, PD169316, RWJ67657, and BIRB796).

Resolvable nanoTAK relaxation traces were obtained for five of the six inhibitors (Figs. 3 and S3). Remarkably, even the smallest +5 °C jump from 20 to 25 °C yielded high-quality data, demonstrating that the method can deliver directly comparable room-temperature kinetics for protein–small molecule systems in addition to its native access to higher, physiologically relevant temperatures. While the association rate *k*_on_ remained relatively constant around values in the 10^6^ to 10^7^ M^−1^s^−1^ range for each of these five inhibitors, the dissociation rate *k*_off_ increased substantially with higher temperatures, driving the decrease in binding affinity.

**Fig. 3:**
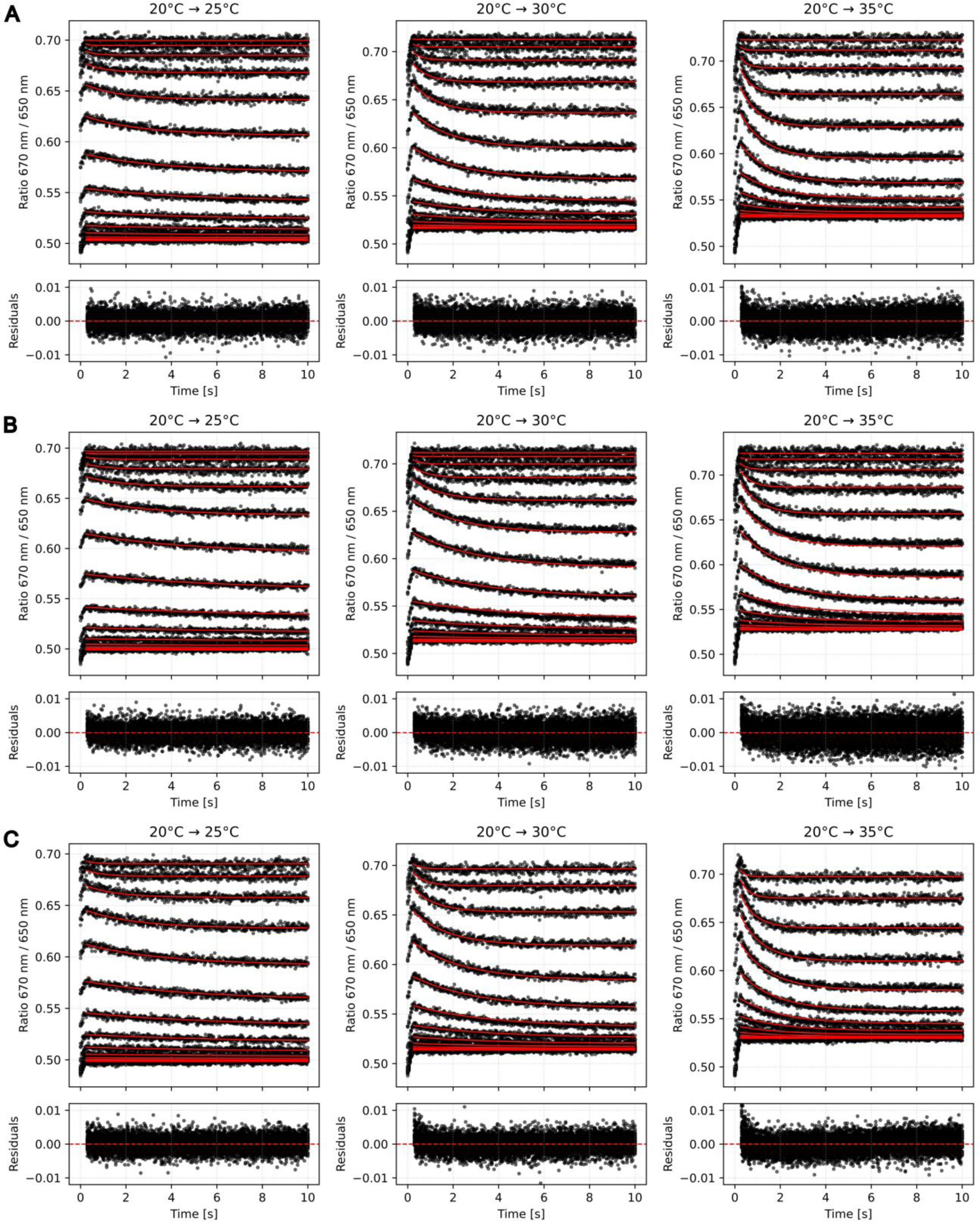
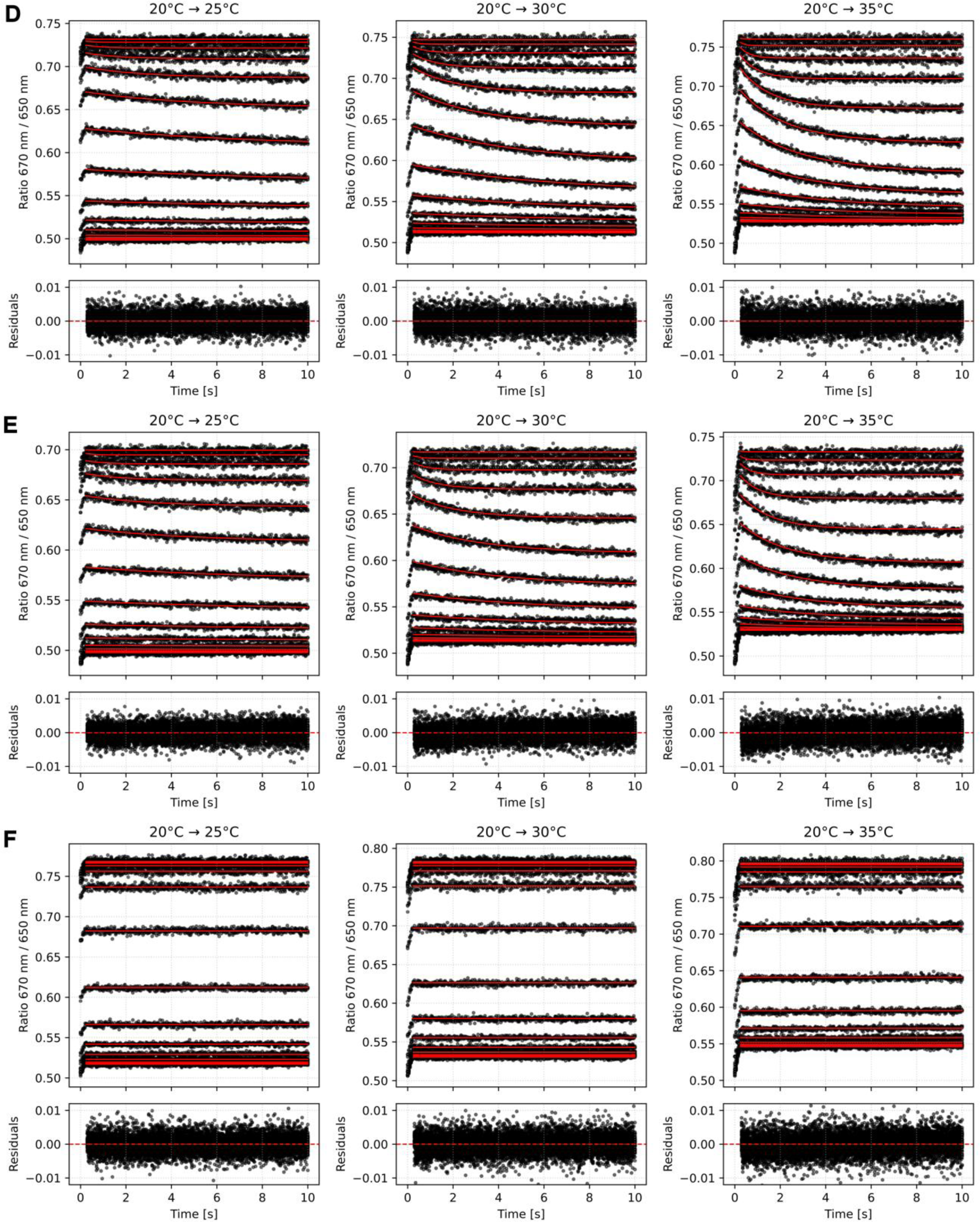
nanoTAK measurements of p38α binding to small molecule inhibitors. SB203580 (**A**), SB202190 (**B**), SB239063 (**C**), RWJ67657 (**D**), PD169316 (**E**), and BIRB796 (**F**). From left to right, the panels show relaxation traces (black) and global fits (red) following temperature jumps of +5, +10, and +15 °C from an initial temperature of 20 °C. The kinetic traces represent merged curves from triplicate measurements. Except for BIRB796, all inhibitors produced clear relaxation kinetics during re-equilibration following the temperature jump.

The one notable exception was BIRB796, which is a picomolar, very-slow-off-rate, allosteric inhibitor^12^. Compared to the other inhibitors, the measured relaxation traces by nanoTAK were flat, because the tight complex did practically not re-equilibrate within the short measurement window of only 10 seconds. Consistent with this interpretation, repeated Spectral Shift measurements over several hours after preparation of the dilution series showed that the apparent dose response curve at 20 °C continued to evolve before reaching a stable response, providing direct evidence of the very slow equilibration kinetics of this interaction (Fig. S4).

Among the tested inhibitors, SB203580 (Fig. 3A) is the best characterized and therefore provides a useful benchmark for comparing nanoTAK with SPR (Table 2). At 25 °C, nanoTAK measured *k*_on_ = 8.2 × 10^6^ M^−1^s^−1^ and *k*_off_ = 0.14 s^−1^, corresponding to a *K*_d_ of 17.3 nM. These values are in good agreement with those reported from SPR at the same temperature, which yielded *K*_d_ = 11.5 nM, *k*_on_ = 1.5 × 10^7^ M^−1^s^−1^ and *k*_off_ = 0.18 s^−1^.^13^ At 35 °C, the same SPR study measured *K*_d_ = 28.9 nM, *k*_on_ = 2.3 × 10^7^ M^−1^s^−1^, and *k*_off_ = 0.62 s^−1^. The greater temperature dependence of *k*_off_ than of *k*_on_ is consistent with the nanoTAK results and supports a temperature-induced weakening of the interaction driven primarily by accelerated dissociation. A second SPR study reported *K*_d_ = 21.7 nM, *k*_on_ = 1.0 × 10^7^ M^−1^s^−1^ and *k*_off_ = 0.22 s^−1^, although the measurement temperature was not specified.^14^ More recent measurements using a high-throughput fluorescence-probe tracer assay are also in close agreement with nanoTAK data. For SB203580, that study reported *k*_on_ = 5.24 × 10^6^ M^−1^s^−1^ and *k*_off_ = 0.106 s^−1^.^15^ Likewise, for another inhibitor SB202190 (Fig. 3B), nanoTAK measured *k*_on_ = 9.42 × 10^6^ M^−1^s^−1^ and *k*_off_ = 0.0799 s^−1^, compared with published values of 1.32 × 10^7^ M^−1^s^−1^ and 0.116 s^−1^, respectively.^15^ The kinetic and thermodynamic results of all molecules can be found in Table S1 of the Supplementary Information. Overall, these comparisons demonstrate that the nanoTAK results are consistent with independent SPR and fluorescence-based measurements and reproduce the expected temperature-dependent weakening of the interaction, primarily through an increase in the dissociation rate.

**Table 2:** Kinetic and thermodynamic parameters of p38α–SB203580 binding.

| Table 2 \| Kinetic and thermodynamic parameters of p38α-SB203580 binding. | | | | | | | |
| --- | --- | --- | --- | --- | --- | --- | --- |
| Method | T (°C) | K<sub>d</sub> (nM) | k<sub>on</sub> (10<sup>6</sup> M<sup>-1</sup>s<sup>-1</sup>) | k<sub>off</sub> (s<sup>-1</sup>) | $\Delta G$ (kJ × mol<sup>-1</sup>) | $\Delta H$ (kJ × mol<sup>-1</sup>) | -T$\Delta S$ (kJ × mol<sup>-1</sup>) |
| SpS | 20 | 11.2 ± 0.3 | — | — | — | — | — |
| nanoTAK | 25 | 17.3 ± 0.5 | 8.2 ± 0.2 | 0.14 ± 0.01 | -44.6 ± 0.1 | -61.7 ± 5.0 | 17.1 ± 5.0 |
| nanoTAK | 30 | 26.8 ± 0.7 | 9.8 ± 0.1 | 0.26 ± 0.01 | -44.6 ± 0.1 | -64.5 ± 2.4 | 19.8 ± 2.4 |
| nanoTAK | 35 | 41.9 ± 1.0 | 11.8 ± 0.1 | 0.49 ± 0.02 | -44.6 ± 0.1 | -65.8 ± 1.6 | 21.2 ± 1.6 |
| SPR<sup>13</sup> | 25 | 11.5 ± 0.8 | 15 ± 3 | 0.18 ± 0.05 | — | — | — |
| SPR<sup>13</sup> | 35 | 28.9 ± 6.1 | 23 ± 5 | 0.62 ± 0.01 | — | — | — |
| SPR<sup>14</sup> | N/A | 21.7 | 10 | 0.22 | — | — | — |
| kPCA<sup>15</sup> | N/A | 66 ± 6 | 5.24 | 0.106 | — | — | — |

In summary, the smaller enthalpic changes of the p38α system compared to DNA hybridization did not limit the nanoTAK method. Although the +5 °C jump only caused small movements of the equilibrium, the data were still high quality and yielded robust kinetic results, while larger jumps of +10 or +15 °C increased the relaxation amplitude.

### Binary and ternary complex interactions

PROTACs (proteolysis-targeting chimeras) are bifunctional molecules that recruit an E3 ubiquitin ligase to a target protein. Their activity is governed by the kinetics of several interactions, including the binary binding of the PROTAC to each protein and the subsequent formation of the ternary complex that brings the two proteins together. Because these interactions involve multiple proteins and may be affected by immobilization, PROTACs provide a compelling test case for an in-solution kinetics approach. Here we studied MZ1, a well-characterized PROTAC that links a BET-bromodomain ligand to a VHL ligand, with the VHL E3-ligase complex (VCB) constituting the labeled target molecule.

The binary VHL–MZ1 interaction is very fast. The relaxation is complete within ∼3 seconds, yet the nanoTAK measurement still resolved it cleanly (Fig. 4A). Three temperature jumps from 20 to 25 °C, 30 °C and 35°C, each performed as three technical replicates on the same capillaries, gave reproducible and internally consistent results (Table 3). The dissociation rate rose monotonically with temperature (*k*_off_ ≈ 0.44 → 0.73 → 1.2 s^−1^) while *k*_on_ stayed near 1 × 10^7^ M^−1^s^−1^, and the determined *K*_d_ at 20 °C was stable across all three series (Fig. S5), confirming that the system returned fully to its starting equilibrium between jumps.

The fitted exothermic enthalpy (Δ*H* ≈ −29.2 to −32.9 kJ mol^−1^) is in excellent agreement with published isothermal titration calorimetry for VCB–MZ1 (Δ*H* ≈ −7.7 kcal mol^−1^ ≈ −32 kJ mol^−1^)^16^. Notably, nanoTAK was able to measure this interaction at and above room temperature, whereas the published SPR data for the same PROTAC was acquired at 12 °C because the interaction was too fast at room temperature^17^.

**Fig. 4:**
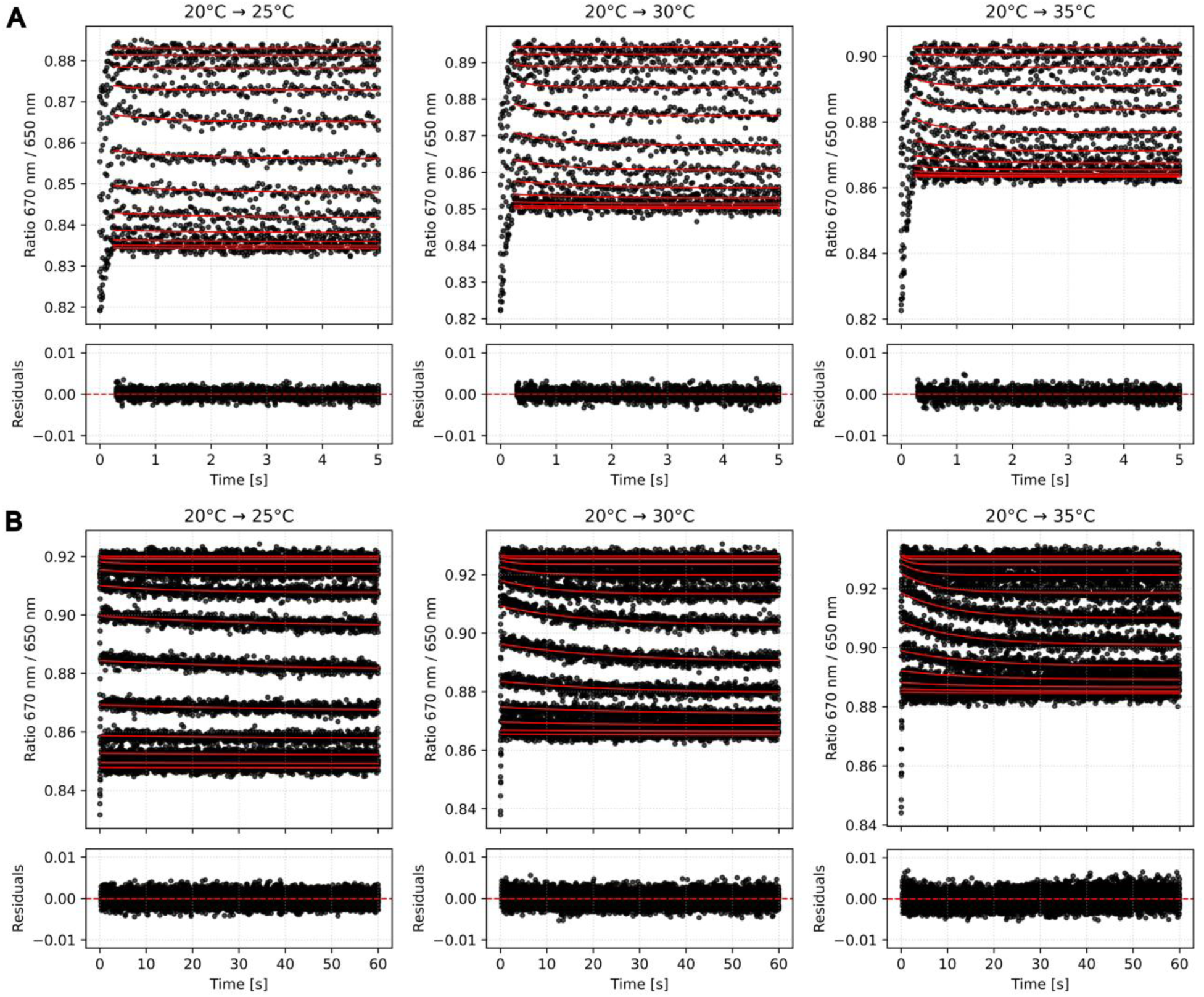
nanoTAK relaxation traces for the VHL–MZ1–BRD4(BD2) PROTAC system. Binary VHL–MZ1. (**A**) and ternary VHL–MZ1–BRD4(BD2) complex (**B**). From left to right, the panels show responses to temperature jumps of +5, +10, and +15 °C from an initial temperature of 20 °C. Black traces display merged triplicate measurements; red curves show the global fits. Measurement times were 5 s per capillary for the binary complex and 60 s per capillary for the slower-dissociating ternary complex.

**Table 3:** Kinetic and thermodynamic parameters of VHL–MZ1–BRD4(BD2) binding.

|  |  |  |  |  |  |  |  |
| --- | --- | --- | --- | --- | --- | --- | --- |
| Method | T (°C) | K<sub>d</sub> (nM) | k<sub>on</sub> (10<sup>6</sup> M<sup>-1</sup>s<sup>-1</sup>) | k<sub>off</sub> (s<sup>-1</sup>) | ΔG (kJ × mol<sup>-1</sup>) | ΔH (kJ × mol<sup>-1</sup>) | -TΔS (kJ × mol<sup>-1</sup>) |
| *Binary complex: VHL-MZ1* |  |  |  |  |  |  |  |
| SpS | 20 | 62.6 ± 4.4 | — | — | — | — | — |
| nanoTAK | 25 | 74.9 ± 5.8 | 5.9 ± 0.7 | 0.44 ± 0.06 | -40.5 ± 0.2 | -29.2 ± 15.6 | -11.3 ± 15.6 |
| nanoTAK | 30 | 96.4 ± 6.3 | 7.6 ± 0.6 | 0.73 ± 0.06 | -40.4 ± 0.2 | -31.0 ± 6.7 | -9.4 ± 6.7 |
| nanoTAK | 35 | 122 ± 9 | 9.3 ± 0.6 | 1.2 ± 0.1 | -40.4 ± 0.2 | -32.9 ± 4.8 | -7.5 ± 4.8 |
| SPR<sup>17</sup> | 12 | 29 | 0.7 | 0.019 | — | — | — |
| ITC<sup>16</sup> | 25 | 66 ± 6 | — | — | -41.1 ± 0.2 | -32.2 ± 0.9 | -8.8 ± 0.9 |
| *Ternary complex: VHL-MZ1, in the presence of 400 nM BRD4(BD2)* |  |  |  |  |  |  |  |
| SpS | 20 | 1.74 ± 0.11 | — | — | — | — | — |
| nanoTAK | 25 | 2.25 ± 0.11 | 4.2 ± 0.3 | 0.0094 ± 0.001 | -49.1 ± 0.2 | -35.6 ± 12.8 | -13.5 ± 12.8 |
| nanoTAK | 30 | 2.97 ± 0.10 | 6.7 ± 0.2 | 0.020 ± 0.001 | -49.2 ± 0.2 | -38.1 ± 5.8 | -11.0 ± 5.8 |
| nanoTAK | 35 | 4.08 ± 0.13 | 9.6 ± 0.3 | 0.039 ± 0.002 | -49.2 ± 0.2 | -44.3 ± 5.3 | -4.9 ± 5.3 |
| SPR<sup>17</sup> | 25 | 1 | 5.9 | 0.006 | — | — | — |
| ITC<sup>16</sup> | 25 | 3.7 ± 0.7 | — | — | -48.1 ± 0.2 | -37.3 ± 0.4 | -10.9 ± 0.8 |
| nanoTAK measurements were performed in triplicate using the same set of capillaries. Thermodynamic parameters for nanoTAK are reported at 20 °C. SpS: Equilibrium Spectral Shift measurements. SPR: surface plasmon resonance. ITC: isothermal titration calorimetry. |  |  |  |  |  |  |  |

We next characterized the ternary VHL–MZ1–BRD4(BD2) complex to investigate how the kinetics of complex formation change when the second protein is present. For this, we reduced the target concentration from 10 nM to 2.5 nM and performed a dilution series of MZ1 in the presence of a constant concentration of 400 nM BRD4(BD2). While the *k*_on_ basically did not change, the off-rate became ∼40-times slower while still rising monotonically with temperature (*k*_off_ ≈ 0.0094 → 0.020 → 0.039 s^−1^, Table 3). Compared to the binary VHL–MZ1 interaction, it is this slower off-rate that accounts for the approximately 30-fold positive cooperativity observed for the ternary complex. The nanoTAK derived kinetic constants at 25 °C (*k*_on_ = 4.2 × 10^6^ M^−1^s^−1^, *k*_off_ = 0.0094 s^−1^) closely match published SPR data (*k*_on_ = 5.9 × 10^6^ M^−1^s^−1^, *k*_off_ = 0.006 s^−1^)^17^. The obtained binding enthalpy for the ternary complex (Δ*H* ≈ −35.6 to −44.3 kJ mol^−1^) is also in excellent agreement with published isothermal titration calorimetry for VCB–MZ1–BRD4(BD2) (Δ*H* ≈ −8.9 kcal mol^−1^ ≈ −37 kJ mol^−1^)^16^.

Taken together, the results with this PROTAC system show that nanoTAK as a single solution-based assay can deliver accurate kinetic and thermodynamic parameters for binary and ternary interactions of a degrader.

## Discussion

Across the three validated systems, nanoTAK measured dissociation rates from 0.009 s^−1^ to just below 2 s^−1^, and association rates from approximately 2 × 10^5^ to 2 × 10^7^ M^−1^s^−1^. Figure 5 summarizes the rate constants determined for the two investigated protein systems in a kinetic rate map, showing how the interactions are distributed across this range and how their rate constants shift with temperature.

**Fig. 5:**
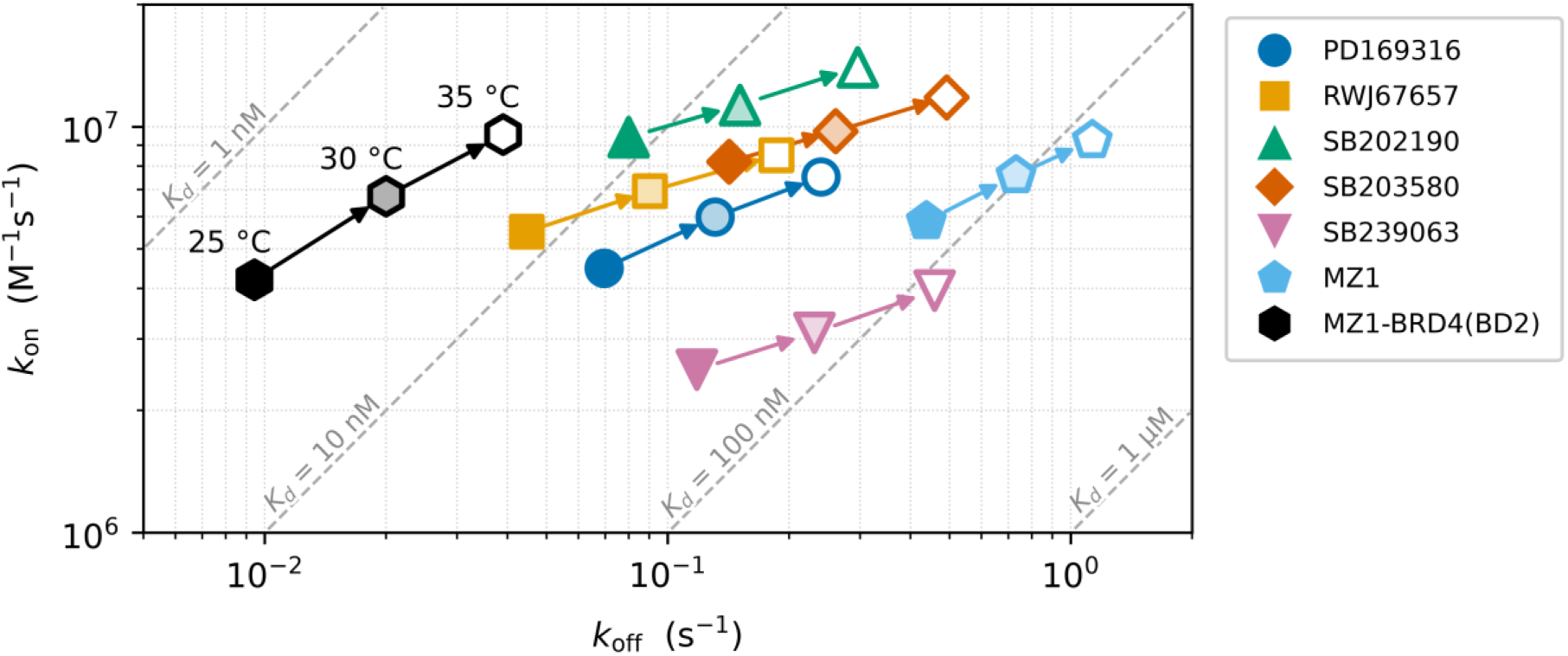
Kinetic rate map of p38α and VHL interactions. The association rate constant (*k*_on_, y-axis) is plotted against the dissociation rate constant (*k*_off_, x-axis) on logarithmic scales for the indicated ligands. Ligands are color-coded: PD169316 (blue circles), RWJ67657 (orange squares), SB202190 (green upward triangles), SB203580 (red diamonds), and SB239063 (purple downward triangles) for p38α, and MZ1 (sky blue pentagons) and MZ1-BRD4(BD2) (black hexagons) for VHL. Symbol fill indicates the measurement temperature: closed symbols, 25 °C; half-filled symbols, 30 °C; open symbols, 35 °C. Arrows connect consecutive temperatures for each ligand, from 25 to 30 °C and from 30 to 35 °C. Dashed diagonal lines represent constant equilibrium dissociation constants (*K*_d_ *= k*_off_ */ k*_on_) of 1 nM, 10 nM, 100 nM, and 1 µM. For all measured interactions, the on-rate slightly increases with temperature, while the off-rate increases more, so overall the binding affinity becomes weaker.

At the fast end of the range, nanoTAK can match or exceed the practical measurement window of surface-based methods such as SPR and BLI. Because the method requires no mixing step and is not subject to mass-transport limitations, it can capture rapid association events without the fluidic constraints that can distort sensorgrams at short timescales. The current limiting factor is the temperature-step rise time of 250 ms. When an interaction already re-equilibrates substantially within this interval, only partial relaxation is observed, which makes reliable rate-constant extraction difficult. Dissociation rates above approximately 2 s^−1^ therefore cannot be quantified precisely with this method. Nevertheless, nanoTAK can still detect a change in *K*_d_ after the temperature jump, even when the kinetics are too fast to resolve, which allows the binding enthalpy to be determined.

At the slow end, very long-lived complexes present another limitation. If *k*_off_ is so small that the complex does not detectably re-equilibrate within the measurement window, relaxation traces appear flat and no kinetic information can be extracted. BIRB796, a picomolar, slow-dissociating inhibitor of p38α, illustrates this limitation: despite a clear equilibrium binding signal across the dilution series, nanoTAK traces were flat at all three temperature jumps tested (Fig. 3F). High-affinity antibody–antigen pairs with sub-nanomolar *K*_d_ values may present a similar challenge. Such interactions are better addressed by mixing-based approaches that monitor association or dissociation over minutes to hours. While the measurement time per capillary can in principle be extended beyond 60 seconds, this reduces throughput due to the sequential scanning architecture and increases the risk of photobleaching of the fluorescent label. In practice, nanoTAK is most reliable for interactions with *k*_off_ ≥ ∼0.005 s^−1^, corresponding to complex residence times up to a few minutes.

Independent from the kinetic constraints, the most stringent condition for nanoTAK is a non-zero binding enthalpy. Since the method is based on the van’t Hoff relationship, only interactions for which *K*_d_ depends on temperature, i.e., enthalpically driven interactions, produce a measurable relaxation signal after a temperature step. A purely entropy-driven interaction would show no shift in the bound fraction upon heating and therefore no resolvable kinetics. In practice, however, most drug-relevant molecular interactions carry at least a partial enthalpic contribution^18–20^. Even systems with modest Δ*H* values, such as the binary VHL–MZ1 interaction (Δ*H* ≈ −30 kJ mol^−1^), yielded clean and analyzable relaxation traces when a larger temperature step (+10 or +15 °C) was applied. Thus, binding kinetics that cannot be resolved with a +5 °C temperature step may become resolvable by simply increasing the jump amplitude.

The thermal stability of the sample at the elevated temperature *T*_1_ is another important point to consider. Re-establishment of the initial equilibrium Spectral Shift signal after each temperature jump serves as a built-in quality control for reversibility: if the system does not return to its pre-jump dose response, irreversible denaturation or aggregation may have occurred, and the kinetic data should be interpreted with caution or discarded. In the systems studied here, all temperature jumps up to +15 °C were fully reversible across repeated measurements on the same set of capillaries. For less thermostable proteins, the accessible jump range may need to be reduced. Operating from a lower starting temperature (e.g., 20 °C instead of 25 °C) can also extend the safety margin before approaching a protein’s unfolding transition.

Finally, the global kinetic model used to fit nanoTAK data assumes a reversible 1:1 binding equilibrium. Systems that deviate from this, including targets with multiple independent or competing binding sites, may yield apparent rate constants that do not correspond cleanly to elementary microscopic rates. The ternary PROTAC system (VHL–MZ1–BRD4(BD2)) demonstrated that cooperative multi-protein assemblies can nevertheless be characterized within this framework by treating the interaction as an effective 1:1 process between the binary complex and the third binding partner, yielding rate constants in close agreement with published SPR data. Taken together, these considerations map out the conditions under which nanoTAK can yield reliable, immobilization-free binding kinetics directly in solution with low sample consumption.

## Conclusion

We have established nanoTAK as a practical, immobilization-free method for measuring in-solution binding kinetics in standard glass capillaries. It requires only microliter sample volumes and the same preparation as a conventional equilibrium binding affinity measurement. A calibrated, spatially homogeneous temperature step displaces an enthalpically driven equilibrium, and global fitting of the resulting relaxation across a dilution series allows the extraction of *k*_on_, *k*_off_, and *K*_d_ at the elevated temperature. Validation on high-enthalpy DNA hybridization confirmed the developed kinetic model under near-ideal conditions. A panel of p38α inhibitors then demonstrated that the approach extends to classical protein–small molecule interactions with much smaller enthalpic changes. Finally, the MZ1 PROTAC with the VHL complex and the bromodomain BRD4(BD2) showed that binary and ternary interactions of a degrader can be characterized side by side in solution, resolving even fast kinetics that can be challenging for surface methods. By measuring association and dissociation rate constants at and above room temperature, nanoTAK offers an alternative for characterizing intermediate-to-fast binding kinetics directly in solution, from room temperature to physiologically relevant temperatures. Temperature-dependent measurements also enable rapid thermodynamic characterization, helping reveal whether an interaction is predominantly enthalpy- or entropy-driven.

## Author Contributions

A.L. conceived and developed the nanoTAK method, developed the global fitting model, designed and supervised the research, and wrote the manuscript. C.O. led the technical development of the nanoTAK instrumentation, designed and implemented the optical measurement system, and devised and implemented the IR-laser calibration procedure. C.O., N.-E.G., and O.R. developed control software for nanoTAK measurements. A.L., P.S., and M.M. labeled proteins and performed nanoTAK measurements. C.O. and P.S. proofread the manuscript. All authors approved the final manuscript.

