## Supplementary Information for "In-Solution Measurements of Binding Kinetics and Thermodynamics by nano Temperature Alteration Kinetics"

**Content:**


- Note S1: Analytical Solution for 1:1 Binding Kinetics in Solution
- Figure S1: nanoTAK measurements of DNA hybridization at a capillary tray temperature of 20 °C.
- Figure S2: nanoTAK measurements of DNA hybridization at a capillary tray temperature of 25 °C.
- Figure S3: Initial equilibrium dose response data for nanoTAK measurements of p38α binding to small molecule inhibitors.
- Figure S4: Time-dependent dose-response curves for p38α binding to BIRB796.
- Figure S5: Initial equilibrium dose response data for nanoTAK measurements of the VHL complex with PROTAC MZ1 and bromodomain BRD4(BD2).
- Table S1: Kinetic and thermodynamic parameters from nanoTAK measurements of p38α.

### Supplementary Note S1

#### Analytical Solution for 1:1 Binding Kinetics in Solution

Quantitative characterization of molecular interactions is central to drug discovery and biophysical characterization. A fundamental quantity in this context is the dissociation constant $K_\mathrm{d}$, which reports on binding affinity, alongside the association and dissociation rate constants ($k_\mathrm{on}$ and $k_\mathrm{off}$), which together describe the kinetics of complex formation and dissociation.

For a reversible 1:1 binding interaction the system can be described by a single ordinary differential equation (ODE). While this ODE is nonlinear, it surprisingly allows an exact closed-form analytical solution. The following derivation presents this analytical solution in full.

#### Mathematical Derivation

We consider reversible binding between a target $\mathrm{T}$ and a ligand $\mathrm{L}$ under a 1:1 stoichiometric model, yielding the complex $\mathrm{TL}$. The free concentrations of target, ligand, and complex are denoted by $[\mathrm{T}]$, $[\mathrm{L}]$, and $[\mathrm{TL}]$, respectively, whereas $[\mathrm{T}]_\mathrm{t}$ and $[\mathrm{L}]_\mathrm{t}$ denote the corresponding total concentration.

Mass conservation requires that the total target and ligand concentrations equal the sum of their free and bound contributions:

$$[\mathrm{L}]_\mathrm{t} = [\mathrm{L}] + [\mathrm{TL}] \quad \rightarrow \quad [\mathrm{L}] = [\mathrm{L}]_\mathrm{t} - [\mathrm{TL}] \tag{S1}$$

$$[\mathrm{T}]_\mathrm{t} = [\mathrm{T}] + [\mathrm{TL}] \quad \rightarrow \quad [\mathrm{T}] = [\mathrm{T}]_\mathrm{t} - [\mathrm{TL}] \tag{S2}$$

The rate of change of the complex concentration is determined by association of free target and ligand with rate constant $k_\mathrm{on}$ and by dissociation of the complex with rate constant $k_\mathrm{off}$:

$$\frac{\mathrm{d}[\mathrm{TL}]}{\mathrm{d}t} = k_\mathrm{on} \cdot [\mathrm{T}] \cdot [\mathrm{L}] - k_\mathrm{off} \cdot [\mathrm{TL}] \tag{S3}$$

At equilibrium, the net rate of complex formation vanishes. The equilibrium condition therefore defines the dissociation constant $K_\mathrm{d}$ as follows:

$$\frac{\mathrm{d[TL]}}{\mathrm{d}t} = 0 \quad \rightarrow \quad \frac{[\mathrm{T}] \cdot [\mathrm{L}]}{[\mathrm{TL}]} = \frac{k_\mathrm{off}}{k_\mathrm{on}} = K_\mathrm{d} \tag{S4}$$

From equations (S1) through (S4), we derive:

$$\begin{aligned}\frac{\mathrm{d[TL]}}{\mathrm{d}t} &= k_\mathrm{on} \cdot ([\mathrm{T}]_\mathrm{t} - [\mathrm{TL}]) \cdot ([\mathrm{L}]_\mathrm{t} - [\mathrm{TL}]) - k_\mathrm{off} \cdot [\mathrm{TL}] \\ &= k_\mathrm{on} \cdot \left([\mathrm{T}]_\mathrm{t} \cdot [\mathrm{L}]_\mathrm{t} - [\mathrm{T}]_\mathrm{t} \cdot [\mathrm{TL}] - [\mathrm{TL}] \cdot [\mathrm{L}]_\mathrm{t} + [\mathrm{TL}]^2 - \frac{k_\mathrm{off}}{k_\mathrm{on}} \cdot [\mathrm{TL}]\right) \\ &= k_\mathrm{on} \cdot ([\mathrm{TL}]^2 - ([\mathrm{T}]_\mathrm{t} + [\mathrm{L}]_\mathrm{t} + K_\mathrm{d}) \cdot [\mathrm{TL}] + [\mathrm{T}]_\mathrm{t} \cdot [\mathrm{L}]_\mathrm{t})\end{aligned} \tag{S5}$$

Introducing the fraction of bound target as $x(t) = [\mathrm{TL}](t)/[\mathrm{T}]_\mathrm{t}$, Eq. (S5) can be rewritten as

$$\frac{\mathrm{d}x}{\mathrm{d}t} = k_\mathrm{on} \cdot [\mathrm{T}]_\mathrm{t} \cdot \left(x^2 - \left(1 + \frac{[\mathrm{L}]_\mathrm{t} + K_\mathrm{d}}{[\mathrm{T}]_\mathrm{t}}\right) \cdot x + \frac{[\mathrm{L}]_\mathrm{t}}{[\mathrm{T}]_\mathrm{t}}\right) \tag{S6}$$

The quadratic polynomial in Eq. (S6) can be factorized as:

$$\frac{\mathrm{d}x}{\mathrm{d}t} = k_\mathrm{on} \cdot [\mathrm{T}]_\mathrm{t} \cdot (x - x_1) \cdot (x - x_2) \tag{S7}$$

where $x_1$ and $\mathrm{x}_2$ are the two roots of the quadratic expression, given by

$$x_{1,2} = \frac{1}{2}\left(1 + \frac{[\mathrm{L}]_\mathrm{t} + K_\mathrm{d}}{[\mathrm{T}]_\mathrm{t}} \pm D\right) \tag{S8}$$

with discriminant

$$D = \sqrt{\left(1 + \frac{[\mathrm{L}]_\mathrm{t} + K_\mathrm{d}}{[\mathrm{T}]_\mathrm{t}}\right)^2 - 4 \cdot \frac{[\mathrm{L}]_\mathrm{t}}{[\mathrm{T}]_\mathrm{t}}} \tag{S9}$$

Since the complex concentration cannot exceed the total concentration of either binding partner, only the smaller of the two roots is physically meaningful. This root therefore defines the equilibrium fraction of bound target $x_{\mathrm{eq}}$:

$$x_1 = \frac{1}{2}\left(1 + \frac{[\mathrm{L}]_\mathrm{t} + K_\mathrm{d}}{[\mathrm{T}]_\mathrm{t}} - D\right) = x_{\mathrm{eq}} \tag{S10}$$

Further, the following useful identity holds:

$$x_2 - x_1 = D \qquad \rightarrow \quad x_2 = x_1 + D \tag{S11}$$

### SOLVING THE NON-LINEAR ODE

Eq. (S7) can be rearranged to

$$k_{\mathrm{on}} \cdot [\mathrm{T}]_\mathrm{t} \cdot \mathrm{d}t = \frac{1}{(x - x_1) \cdot (x - x_2)} \cdot \mathrm{d}x \tag{S12}$$

Using partial-fraction decomposition, the fraction in Eq. (S12) can be rewritten as

$$\begin{aligned} \frac{1}{(x - x_1) \cdot (x - x_2)} = \frac{1}{x_2 - x_1} \cdot \frac{x_2 - x_1}{(x - x_1) \cdot (\mathrm{x} - x_2)} = \frac{1}{x_2 - x_1} \cdot \frac{(x - x_1) - (x - x_2)}{(x - x_1) \cdot (\mathrm{x} - x_2)} \\ = \frac{1}{x_2 - x_1} \cdot \left(\frac{1}{x - x_2} - \frac{1}{x - x_1}\right) = \frac{1}{D} \cdot \left(\frac{1}{x - x_2} - \frac{1}{x - x_1}\right) \end{aligned} \tag{S13}$$

Substituting $k = D \cdot k_{\mathrm{on}} \cdot [\mathrm{T}]_\mathrm{t}$ we can further simplify to

$$k \cdot \mathrm{d}t = \frac{1}{x - x_2} \cdot \mathrm{d}x - \frac{1}{x - x_1} \cdot \mathrm{d}x \tag{S14}$$

Finally, integration with respect to time yields

$$k \cdot t + C = \ln|\mathrm{x} - x_2| - \ln|x - x_1| = \ln\left|\frac{x - x_2}{x - x_1}\right| \tag{S15}$$

where $C$ is an integration constant determined by the initial condition.

### Initial condition

Assuming that the initial fraction bound at $t = 0$ is $x_0$, Eq. (S15) implies

$$C = \ln\left|\frac{x_0 - x_2}{x_0 - x_1}\right| \tag{S16}$$

and therefore

$$k \cdot t = -\ln\left|\frac{x_0 - x_2}{x_0 - x_1}\right| + \ln\left|\frac{x - x_2}{x - x_1}\right| \tag{S17}$$

Exponentiating both sides of Eq. (S17) gives

$$\mathrm{e}^{kt} = \left|\frac{x_0 - x_1}{x_0 - x_2} \cdot \frac{x - x_2}{x - x_1}\right| \tag{S18}$$

Since $\mathrm{x}(t) < x_2$ for all $t$, both factors in Eq. (S18) containing $x_2$ are negative. Moreover, as $\mathrm{x}(t)$ approaches $x_{\mathrm{eq}} = x_1$ monotonically without crossing it, the factors containing $x_1$ have the same sign in numerator and denominator. Hence, the expression is always positive, and Eq. (S18) may be written without absolute values as

$$\mathrm{e}^{kt} = \frac{x_0 - x_1}{x_0 - x_2} \cdot \frac{x - x_2}{x - x_1} = A \cdot \frac{x - x_2}{x - x_1} \tag{S19}$$

with

$$A = \frac{x_0 - x_1}{x_0 - x_2} = \frac{x_0 - x_{\mathrm{eq}}}{x_0 - x_{\mathrm{eq}} - D} = \frac{x_{\mathrm{eq}} - x_0}{D + x_{\mathrm{eq}} - x_0} \tag{S20}$$

Eq. (S19) can be rearranged to:

$$\mathrm{e}^{kt} \cdot x - \mathrm{e}^{kt} \cdot x_1 = A \cdot x - A \cdot x_2 \quad \rightarrow \quad \mathrm{e}^{kt} \cdot x - A \cdot x = \mathrm{e}^{kt} \cdot x_1 - A \cdot x_2 \tag{S21}$$

Solving Eq. (S21) for $x$ finally yields:

$$x(t) = \frac{\mathrm{e}^{kt} \cdot x_1 - A \cdot x_2}{\mathrm{e}^{kt} - A} = \frac{\mathrm{e}^{kt} \cdot x_{\mathrm{eq}} - A \cdot \left(x_{\mathrm{eq}} + D\right)}{\mathrm{e}^{kt} - A} = x_{\mathrm{eq}} - D \cdot \frac{A}{\mathrm{e}^{kt} - A} \tag{S22}$$

For improved numerical stability, Eq. (S22) can also be rewritten as:

$$x(t) = x_{\mathrm{eq}} - D \cdot \frac{A \cdot \mathrm{e}^{-kt}}{1 - A \cdot \mathrm{e}^{-kt}} \tag{S23}$$

which is identical to Eq. (2) of the main text.

#### Special Cases

For the special cases considered below, it is convenient to first evaluate the inverse of $A$:

$$A^{-1} = \frac{D + x_{\mathrm{eq}} - x_0}{x_{\mathrm{eq}} - x_0} = 1 + \frac{D}{x_{\mathrm{eq}} - x_0} \tag{S24}$$

The trivial case $x_{\mathrm{eq}} = x_0$ is excluded here.

Using this expression, Eq. (S22) can be rewritten as

$$x(t) = x_{\mathrm{eq}} - \frac{D}{A^{-1} \cdot \mathrm{e}^{kt} - 1} \tag{S25}$$

**Initial condition:** $t = 0$

At the initial time point, the solution correctly reproduces the starting fraction bound, $x_0$:

$$x(t = 0) = x_{\mathrm{eq}} - \frac{D}{A^{-1} - 1} = x_{\mathrm{eq}} - \frac{D \cdot (x_{\mathrm{eq}} - x_0)}{D} = x_0 \tag{S26}$$

This confirms that the expression satisfies the required initial condition.

**Long-time limit:** $t \to \infty$

In the long-time limit, the exponential term in the denominator grows without bound, causing the second term to vanish. As a result, the solution approaches the equilibrium fraction bound, $x_{\mathrm{eq}}$:

$$x(t \to \infty) = \lim_{t \to \infty} \left( x_{\mathrm{eq}} - \frac{D}{A^{-1} \cdot \mathrm{e}^{kt} - 1} \right) = x_{\mathrm{eq}} \tag{S27}$$

**Vanishing target concentration limit:** $[\mathrm{T}]_\mathrm{t} \to 0$

In the limit of vanishing total target concentration, ligand depletion becomes negligible, and the solution is expected to reduce to the familiar pseudo-first-order form.

To show this, we first consider the discriminant $D$. As $[\mathrm{T}]_\mathrm{t} \to 0$,

$$D = \sqrt{\left(1 + \frac{[\mathrm{L}]_\mathrm{t} + K_\mathrm{d}}{[\mathrm{T}]_\mathrm{t}}\right)^2 - 4 \cdot \frac{[\mathrm{L}]_\mathrm{t}}{[\mathrm{T}]_\mathrm{t}}} \approx \frac{[\mathrm{L}]_\mathrm{t} + K_\mathrm{d}}{[\mathrm{T}]_\mathrm{t}} \tag{S28}$$

Using this result, the kinetic rate constant $k$ simplifies to

$$k = \frac{[\mathrm{L}]_\mathrm{t} + K_\mathrm{d}}{[\mathrm{T}]_\mathrm{t}} \cdot k_\mathrm{on} \cdot [\mathrm{T}]_\mathrm{t} = ([\mathrm{L}]_\mathrm{t} + K_\mathrm{d}) \cdot k_\mathrm{on} = [\mathrm{L}]_\mathrm{t} \cdot k_\mathrm{on} + k_\mathrm{off} \tag{S29}$$

Finally, in the limit $D \to \infty$, we obtain

$$x = x_\mathrm{eq} - \frac{D}{A^{-1} \cdot \mathrm{e}^{kt} - 1} = x_\mathrm{eq} - \frac{D}{\left(1 + \frac{D}{x_\mathrm{eq} - x_0}\right) \cdot \mathrm{e}^{kt} - 1} \approx x_\mathrm{eq} - \frac{D}{\frac{\mathrm{D}}{x_\mathrm{eq} - x_0} \cdot \mathrm{e}^{kt}} \tag{S30}$$

which further simplifies to the familiar form

$$x = x_\mathrm{eq} - \left(x_\mathrm{eq} - x_0\right) \cdot \mathrm{e}^{-kt} \tag{S31}$$

**Non-zero starting fraction bound:** $x_0 \neq 0$

In experiments in which a pre-established binding equilibrium is perturbed by a temperature jump (nanoTAK), the observed in-solution kinetics describe the relaxation toward a new equilibrium state. In this case, the initial fraction bound, $x_0$, can be obtained from the standard 1:1 tight-binding equilibrium expression with initial dissociation constant $K_\mathrm{d}(T_0)$:

$$x_0 = \frac{[\mathrm{L}]_\mathrm{t} + [\mathrm{T}]_\mathrm{t} + K_\mathrm{d}(T_0) - \sqrt{\left([\mathrm{L}]_\mathrm{t} + [\mathrm{T}]_\mathrm{t} + K_\mathrm{d}(T_0)\right)^2 - 4[\mathrm{L}]_\mathrm{t} \cdot [\mathrm{T}]_\mathrm{t}}}{2 \cdot [\mathrm{T}]_\mathrm{t}} \tag{S32}$$

This is Eq. (1) of the main text.

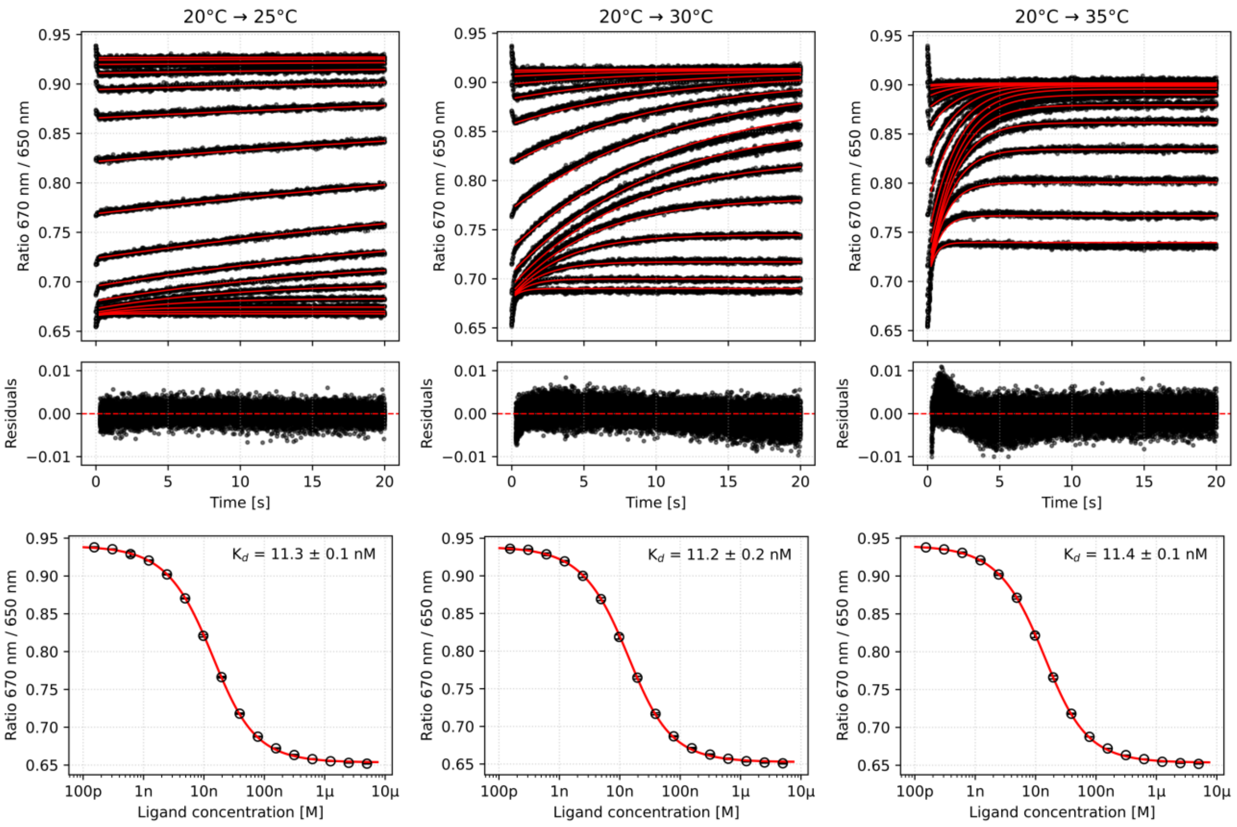


**Fig. S1 | nanoTAK measurements of DNA hybridization at a capillary tray temperature of 20 °C.** A 10-mer duplex containing a Cy5-labeled strand was measured after temperature jumps of +5, +10, and +15 °C, shown from left to right. **Top row:** Experimental relaxation traces (black) and corresponding global in-solution model fits (red). The traces represent triplicate measurements across the dilution series. **Middle row:** Residuals of the global fits. **Bottom row:** Equilibrium dose response data measured before each temperature jump, together with the corresponding fits. The fitted $K_d$ values were consistent across the three measurements, supporting reproducible re-equilibration of the system before each temperature-jump experiment.

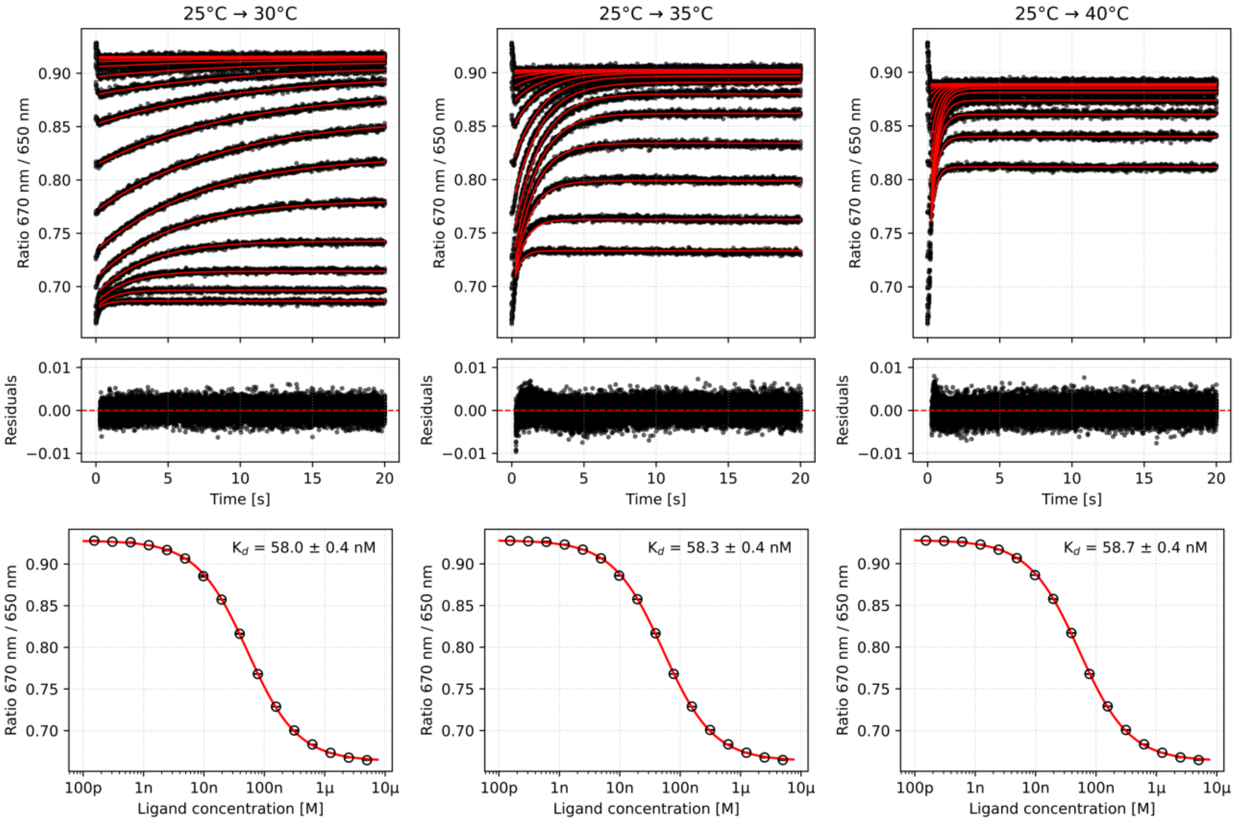


**Fig. S2 | nanoTAK measurements of DNA hybridization at a capillary tray temperature of 25 °C.** A 10-mer duplex containing a Cy5-labeled strand was measured after temperature jumps of +5, +10, and +15 °C, shown from left to right. **Top row:** Experimental relaxation traces (black) and corresponding global in-solution model fits (red). The traces represent triplicate measurements across the dilution series. **Middle row:** Residuals of the global fits. **Bottom row:** Equilibrium dose response data measured before each temperature jump, together with the corresponding fits. The fitted $K_d$ values were consistent across the three measurements, supporting reproducible re-equilibration of the system before each temperature-jump experiment.

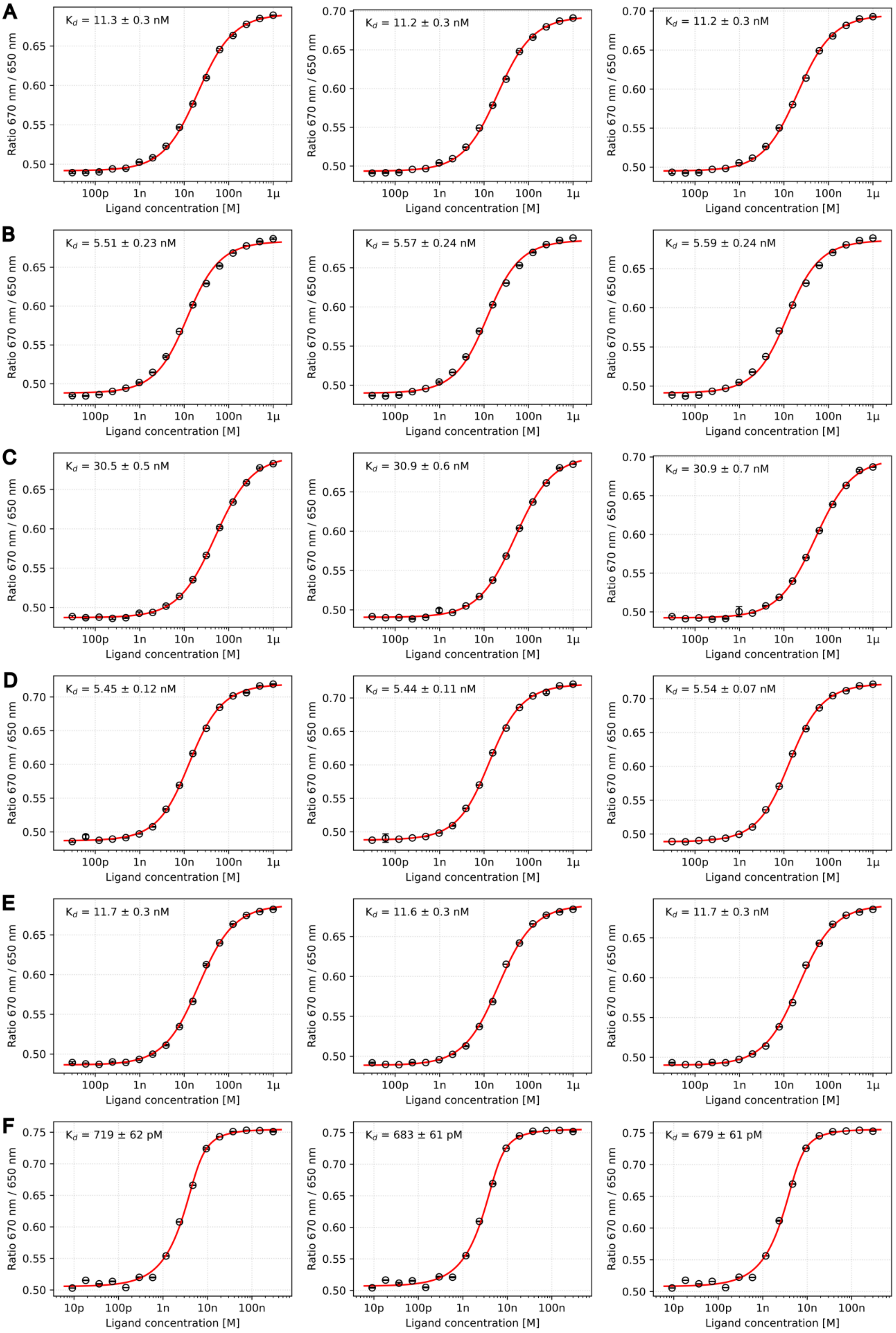
A
$K_d$ = 11.3 ± 0.3 nM
$K_d$ = 11.2 ± 0.3 nM
$K_d$ = 11.2 ± 0.3 nM
B
$K_d$ = 5.51 ± 0.23 nM
$K_d$ = 5.57 ± 0.24 nM
$K_d$ = 5.59 ± 0.24 nM
C
$K_d$ = 30.5 ± 0.5 nM
$K_d$ = 30.9 ± 0.6 nM
$K_d$ = 30.9 ± 0.7 nM
D
$K_d$ = 5.45 ± 0.12 nM
$K_d$ = 5.44 ± 0.11 nM
$K_d$ = 5.54 ± 0.07 nM
E
$K_d$ = 11.7 ± 0.3 nM
$K_d$ = 11.6 ± 0.3 nM
$K_d$ = 11.7 ± 0.3 nM
F
$K_d$ = 719 ± 62 pM
$K_d$ = 683 ± 61 pM
$K_d$ = 679 ± 61 pM
Ratio 670 nm / 650 nm
Ligand concentration [M]

**Fig. S3 | Initial equilibrium dose response data for nanoTAK measurements of p38α binding to small molecule inhibitors.** SB203580 (**A**), SB202190 (**B**), SB239063 (**C**), RWJ67657 (**D**), PD169316 (**E**), and BIRB796 (**F**). Equilibrium dose response data measured before each temperature jump (+5 °C, +10 °C, +15 °C, from left to right), together with the corresponding fits. The consistency of fitted $K_d$ values across the three measurements confirms reproducible re-equilibration of the system before each temperature-jump experiment.

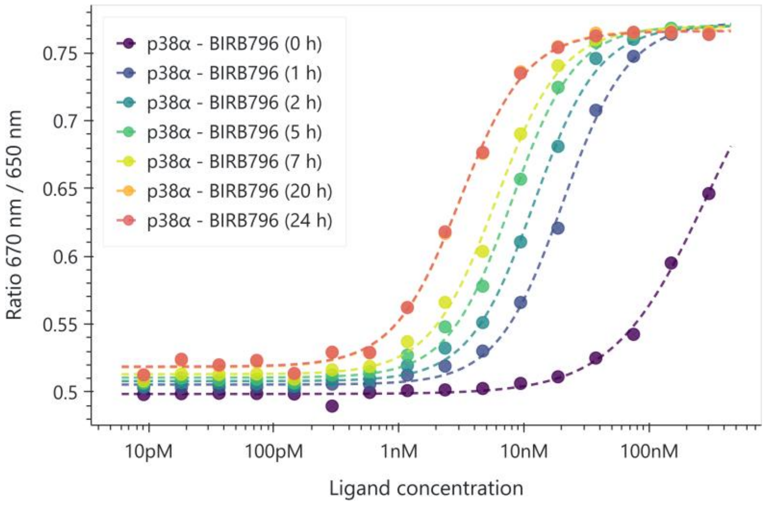


**Fig. S4 | Time-dependent dose-response measurements for p38α binding to BIRB796.** Data were collected immediately after loading the dilution series into the Monolith Omni device (0 h) and again after 1, 2, 5, 7, 20, and 24 h of incubation in the device. Capillaries were sealed to prevent evaporation. The binding data were fitted with a Hill equation. The $EC_{50}$ decreased from 297 nM at 0 h to 3.1 nM at 20 h. The curves were unchanged between 20 and 24 h, indicating that equilibrium was reached by 20 h and that sufficient incubation is required before measuring an equilibrium binding affinity.

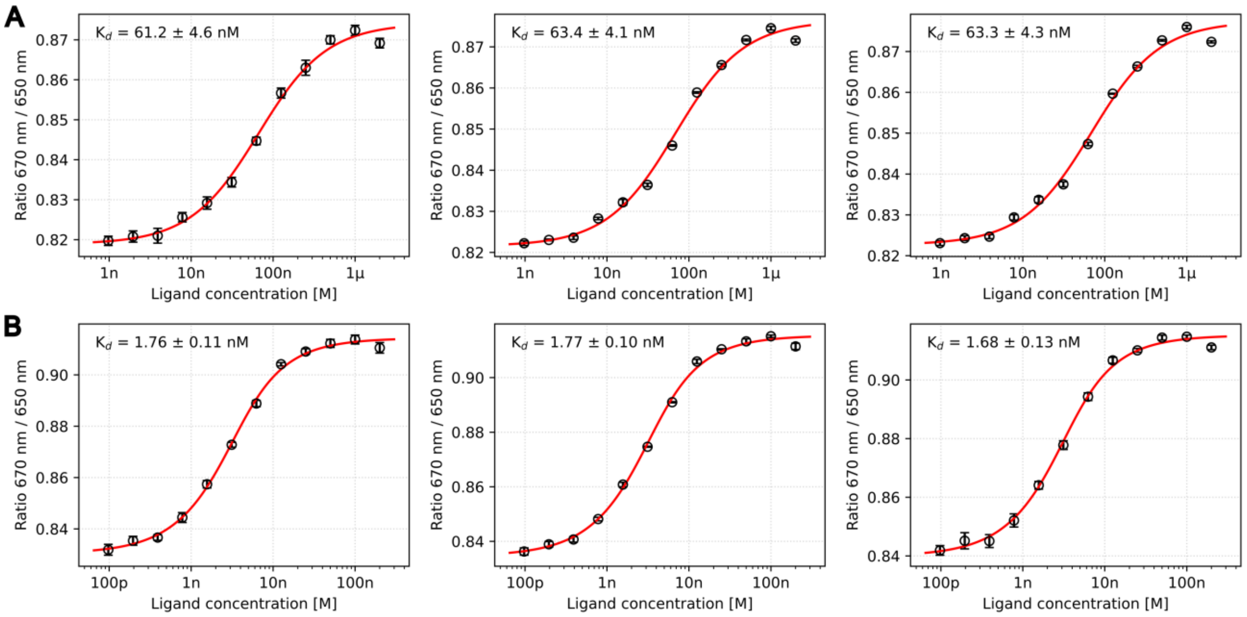


**Fig. S5 | Initial equilibrium dose response data for nanoTAK measurements of the VHL complex with PROTAC MZ1 and bromodomain BRD4(BD2).** The binary complex (**A**) and ternary complex (**B**) were measured at equilibrium before each temperature jump (+5, +10, and +15 °C, from left to right), with the corresponding fits shown. The apparent $K_d$ values indicate increased affinity upon ternary complex formation, decreasing from 61.2–63.3 nM for the binary complex to 1.68–1.77 nM for the ternary complex. The consistent fitted $K_d$ values across the three measurements support reproducible re-equilibration before each temperature-jump experiment.

Table S1 | Kinetic and thermodynamic parameters from nanoTAK measurements of p38α.

| Ligand | Method | T (°C) | $K_d$ (nM) | $k_{on}$ ($10^6$ $M^{-1}s^{-1}$) | $k_{off}$ ($s^{-1}$) | ΔG (kJ × $mol^{-1}$) | ΔH (kJ × $mol^{-1}$) | –TΔS (kJ × $mol^{-1}$) |
|---|---|---|---|---|---|---|---|---|
| SB203580 | SpS | 20 | 11.2 ± 0.3 | — | — | — | — | — |
| | nanoTAK | 25 | 17.3 ± 0.5 | 8.2 ± 0.2 | 0.14 ±0.01 | –44.6 ± 0.1 | –61.7 ± 5.0 | 17.1 ± 5.0 |
| | nanoTAK | 30 | 26.8 ± 0.7 | 9.8 ± 0.1 | 0.26 ± 0.01 | –44.6 ± 0.1 | –64.5 ± 2.4 | 19.8 ± 2.4 |
| | nanoTAK | 35 | 41.9 ± 1.0 | 11.8 ± 0.1 | 0.49 ± 0.02 | –44.6 ± 0.1 | –65.8 ± 1.6 | 21.2 ± 1.6 |
| SB202190 | SpS | 20 | 5.6 ± 0.2 | — | — | — | — | — |
| | nanoTAK | 25 | 8.5 ± 0.4 | 9.4 ± 0.2 | 0.080 ± 0.003 | –46.3 ± 0.1 | –62.7 ± 8.8 | 16.4 ± 8.8 |
| | nanoTAK | 30 | 13.4 ± 0.6 | 11.3 ± 0.2 | 0.15 ± 0.01 | –46.3 ± 0.1 | –64.7 ± 4.5 | 18.3 ± 4.5 |
| | nanoTAK | 35 | 21.2 ± 1.0 | 14.0 ± 0.2 | 0.30 ± 0.02 | –46.3 ± 0.1 | –66.7 ± 3.1 | 20.4 ± 3.1 |
| SB239063 | SpS | 20 | 30.8 ± 0.6 | — | — | — | — | — |
| | nanoTAK | 25 | 46.9 ± 0.79 | 2.5 ± 0.1 | 0.12 ± 0.01 | –42.2 ± 0.1 | –62.5 ± 3.4 | 20.4 ± 3.4 |
| | nanoTAK | 30 | 74 ± 1.5 | 3.1 ± 0.1 | 0.23 ± 0.01 | –42.1 ± 0.1 | –64.5 ± 2.1 | 22.3 ± 2.1 |
| | nanoTAK | 35 | 116 ± 3 | 4.0 ± 0.1 | 0.46 ± 0.02 | –42.1 ± 0.1 | –66.3 ± 1.7 | 24.1 ± 1.7 |
| PD169316 | SpS | 20 | 11.7 ± 0.3 | — | — | — | — | — |
| | nanoTAK | 25 | 15.5 ± 0.4 | 4.5 ± 0.2 | 0.070 ± 0.003 | –44.5 ± 0.1 | –41.7 ± 4.8 | –2.8 ± 4.8 |
| | nanoTAK | 30 | 21.9 ± 0.5 | 6.0 ± 0.2 | 0.13 ± 0.01 | –44.5 ± 0.1 | –47.2 ± 2.5 | 2.7 ± 2.5 |
| | nanoTAK | 35 | 32 ± 0.8 | 7.5 ± 0.2 | 0.24 ± 0.01 | –44.5 ± 0.1 | –50.2 ± 1.6 | 5.7 ± 1.7 |
| RWJ67657 | SpS | 20 | 5.5 ± 0.1 | — | — | — | — | — |
| | nanoTAK | 25 | 8.1 ± 0.2 | 5.5 ± 0.1 | 0.045 ± 0.002 | –46.4 ± 0.1 | –57.8 ± 4.5 | 11.4 ± 4.5 |
| | nanoTAK | 30 | 13 ± 0.3 | 6.9 ± 0.1 | 0.090 ± 0.002 | –46.4 ± 0.1 | –64.4 ± 2.1 | 18 ± 2.1 |
| | nanoTAK | 35 | 21.8 ± 0.3 | 8.5 ± 0.1 | 0.19 ± 0.01 | –46.3 ± 0.1 | –68.6 ± 0.9 | 22.3 ± 0.9 |

Kinetic data were acquired by triplicate measurements on the same set of capillaries. Thermodynamic parameters for nanoTAK are reported at 20 °C. SpS: Equilibrium Spectral Shift measurement.
